# scPDA: Denoising Protein Expression in Droplet-Based Single-Cell Data

**DOI:** 10.1101/2024.12.07.627329

**Authors:** Ouyang Zhu, Jun Li

## Abstract

Droplet-based profiling techniques such as CITE-seq measure the surface protein abundance of single cells, providing crucial information for cell-type identification. However, these measurements are often significantly contaminated by technical noise, which lowers the efficiency of using the gating strategy to identify cell types. Current computational denoising methods have serious limitations, including a strong reliance on often-unavailable empty droplets or null controls, insufficient efficiency due to the ignoring of protein-protein interactions, and a heavy computational load. Here, we introduce scPDA, a new probabilistic model that employs a variational autoencoder to achieve high computational efficiency. scPDA completely eliminates the use of empty droplets, and it shares information across proteins to increase denoising efficiency. Compared to currently available methods, scPDA has removed noise much more thoroughly while preserving biological signals, and it has substantially improved the efficiency of gating-strategy-based cell-type identification, marking a clear advancement in the computational denoising of the protein modality.

## Introduction

The field of single-cell biology has undergone significant development with the emergence of multi-omic sequencing technologies. By utilizing DNA-tagged and oligo-conjugated antibodies, droplet-based single-cell profiling techniques [1, 2], such as CITE-seq [3], ASAP-seq [4], and TEA-seq [5], can now unravel surface protein abundance along with gene expression, providing a more comprehensive understanding of cellular heterogeneity. However, the protein modality of this multi-omic data presents unique challenges in data analysis.

Unlike count data from the mRNA modality, which exhibits high sparsity and is often modeled by a zero-inflated negative binomial distribution [6, 7, 8], count data from the protein modality show a deficiency of zeros: even for genes that are not expressed, their surface protein measurements are not zero. For example, a study on single-cell multi-modality integration [9] reported merely 3,703 zero counts (0.5%) across a panel of 25 antibodies in 30,672 bone marrow cells. Such a low proportion of zero protein counts is typical in droplet-based single-cell protein datasets [10, 11, 12, 13, 14].

The fake non-zero measurements are due to strong background noise caused by both ambient molecules and non-specific bindings [15, 16, 17, 18]. This added background noise poses obstacles for cell-type identification using cell-type-specific protein markers. Consider the same study [9] as an example. While CD4 and CD8 are typically viewed as mutually exclusive markers for helper-T cells and cytotoxic-T cells, respectively, 75% of helper-T cells have CD8 counts greater than 21 in the data, and 75% of cytotoxic-T cells exhibit CD4 counts beyond 15 in the data. This makes using CD4 and CD8 to differentiate the two cell types error-prone. Thus, it is crucial to remove background noise before using data from the protein modality, which is often done by a computational procedure called “denoising.”

Several denoising methods have been developed. Below, we discuss four representative methods, highlighting concepts and considerations essential for understanding our approach. We also comprehensively discuss their limitations. Some of these limitations arise from the design of the methods, while others were identified by us during our extensive use and testing of these packages, which should be considered a contribution of our paper.

An intuitive and widely adopted approach is fitting a two-component Gaussian Mixture Model (GMM) [19] to the expression of each protein across all cells, expecting that the low peak corresponds to noise and the high peak corresponds to the true signal. For each cell, the probability of belonging to the low peak is calculated according to the GMM model, and the cell is classified as noise if this probability exceeds a threshold.

There are several limitations to this approach. First, the GMM cannot adequately characterize sequencing data, which consists of numbers converted from read counts. Although normalization and log-transformation make these data resemble continuous and real-valued data more closely, the processed data still exhibit unequal tails and multiple artificial peaks at the lower end. Second, this method is applied independently to each protein, lacking the ability to share information between proteins (i.e., it ignores protein-protein dependence), thereby limiting its efficiency. Additionally, proteins that do not express in any of the cells, referred to as “non-expressive” proteins [20, 21], exhibit a unimodal expression pattern. In such cases, the bimodal GMM does not apply properly and will likely mistakenly categorize a substantial portion of noise as biological signal (see section S1 of Supplementary Material for details).

A second method, DSB [16], estimates the noise strength and variation by averaging the protein expression in “empty droplets,” which are microfluidic droplets that do not contain cells but still have non-zero counts as they capture ambient molecules or unbound antibodies. DSB then computes standardized data by removing the estimated noise strength and dividing by the estimated noise standard deviation. As an optional additional step, DSB estimates and regresses out the technical component in the protein expression of each cell in order to remove cell-to-cell variation.

A problem with DSB is its reliance on empty droplets. A typical droplet experiment often generates a large number of empty droplets (roughly about ten times the number of cell-containing droplets, the data of interest), but these empty droplets are usually considered useless and thus discarded without being submitted to public databases to save on storage. As a result, most datasets in public databases do not contain empty droplets.

Moreover, even when both empty droplets and other droplets are present in a dataset, accurately identifying empty droplets remains challenging. Typically, empty droplets are identified as droplets whose total counts in the mRNA modality are too small; however, a recent study [18] has demonstrated that only a portion of such droplets are truly empty and suitable for denoising. The identification of this proportion depends on a specific cutoff, and we observe that DSB’s performance is highly sensitive to the choice of this cutoff (details are provided in section S2 of Supplementary Material).

Finally, it is worth mentioning that while the default approach of DSB requires the availability and identification of empty droplets, it has a backup approach—although less accurate— when empty droplets are not available. In such cases, DSB relies on a two-component GMM model and uses the estimated mean and standard deviation of the lower component as those of the noise. This backup approach, apparently, suffers from the limitations of the GMM-based approach, which we have discussed.

A third method is scAR [17]. scAR models *X*_*ij*_, the count of protein *j* in cell *i*, by a binomial distribution Binom(*d*_*i*_, *p*_*ij*_), where *d*_*i*_ is the library size of cell *i*, and

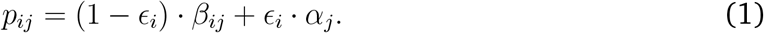

Here, *α*_*j*_ and *β*_*ij*_ are the frequencies for the ambient noise and biological signal, respectively, and *ϵ*_*i*_ is a weight between 0 and 1. While *β*_*ij*_ and *ϵ*_*i*_ are learned from a deep neural network model, *α*_*j*_ is obtained from empty droplets. Thus, scAR has the limitations inherent from the reliance on empty droplets. In cases where empty droplets are unavailable, scAR provides a remedy by estimating *α*_*j*_ through averaging the protein proportion across the cell-containing droplets. However, we have found that *α*_*j*_ estimated this way differs considerably from that estimated from empty droplets (details are provided in section S3 of Supplementary Material), potentially degrading the efficiency of denoising.

A fourth method, DecontPro [18], models *X*_*ij*_ using a Poisson distribution with mean *d*_*i*_ · *p*_*ij*_, where

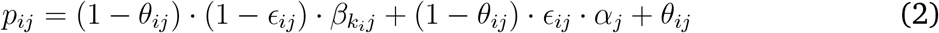

Compared with the *p*_*ij*_ in scAR, which describes only the ambient noise and biological signal, DecontPro further introduces a term *θ*_*ij*_ to describe noise from non-ambient sources. Moreover, unlike scAR, whose biological signal term *β*_*ij*_ is cell-specific, the biological signal term 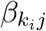 in DecontPro depends only on the cell type *k*_*i*_ to which cell *i* belongs. That is, it assumes that cells from the same cell type have exactly the same expression.

DecontPro also comes with several limitations. First, similar to scAR, DecontPro estimates *α* by averaging either empty droplets or cell-containing droplets, depending on whether empty droplets are available, and consequently, it encounters the same challenges as scAR. Second, DecontPro suffers from poor computational efficiency because it uses computationally demanding variational inference to estimate parameters other than *α*. Third, DecontPro requires the input of a cell-type label *k*_*i*_, which is obtained from clustering the expression of cells before denoising and is not updated after denoising. Thus, the denoising does not enhance the accuracy of cell-type identification. Finally, the assumption that cells from the same cell type have exactly the same expression overlooks within-cell-type variation, which not only exists but can also be highly relevant for many cell types. Ignoring this variation not only impedes understanding of continuous cellular development but also leads to artifacts where all cell clusters appear individually tight and mutually highly distinct.

To overcome the limitations of existing methods, we developed a new method called single-cell protein denoise autoencoder, or scPDA for short. scPDA uses a two-component negative-binomial mixture distribution to model the protein expression data in the form of the original count, instead of the log(1+x) transformed data. Importantly, scPDA addresses the challenge of “non-expressive” proteins by introducing a penalty term to amplify the distinction between the biological signal and the background noise. The parameters of this model are learned through a variational autoencoder network, which is not only computationally efficient but also highly accurate because of its ability to share information across all proteins. Finally, scPDA completely eliminates the use of empty droplets, making it much more applicable to the vast majority of datasets. To demonstrate its efficacy and versatility, we apply scPDA to real datasets that use different technologies to generate the protein modality and compare its performance with other protein denoising methods. The results demonstrate that scPDA significantly outperforms competing methods across these diverse datasets.

## Results

### scPDA removes noise in unstained cells and non-expressive proteins

We will use a CITE-seq dataset provided in [20] and utilized by the DSB paper [16], which includes 53,201 peripheral blood mononuclear cells (PBMCs) on an 87-antibody panel. This dataset is particularly suitable for evaluating denoising methods, as it does not only contain empty droplets but also features a unique aspect: it includes a set of “unstained cells,” which are spike-in cells not expected to bind with any antibodies. These cells serve as groundtruth negative controls, as any non-zero counts from these cells are completely derived from background noise. Moreover, this dataset contains multiple non-expressive proteins (e.g., CD117, CD137, CD138, etc.), which target absent cell types and thus should exhibit zero expression across all cell types in the data. These non-expressive proteins also serve as ground-truth negative controls. Based on these unstained cells and non-expressive proteins, we will be able to evaluate how well denoising methods remove noise: ideally, the expression of any protein in unstained cells and the expression of non-expressive proteins in any cell should be zero after denoising.

Before denoising, positive counts are present across all proteins in unstained cells, confirming the widespread existence of background noise in this data. To exemplify this issue, the left panels in each of Figures 1a-f show the expression of several marker genes in their corresponding cell types (i.e., the cell types for which they are marker genes), and in the unstained cells, in blue and gold colors, respectively. The expression of these genes is expected to be high in the corresponding cell types and zero in the unstained cells. However, as shown in the figures, although on average the blue peaks are higher than the gold, the gold peaks are still non-negligible. These gold peaks are especially significant when comparing across proteins. For example, the average expression of CD8 (shown in Figure 1a) is 35.89 in unstained cells, even higher than the average expression of CD127 (not shown) in its corresponding cell type (i.e., T cells, which have CD127 as one of their marker proteins). Figures 1g-i display the distribution of counts for several non-expressive proteins. All the counts should be zero, which, again, is not the case. Collectively, the widespread positive counts in both proteins in unstained cells and non-expressive proteins underscore the imperative need for denoising.

**Figure 1:**
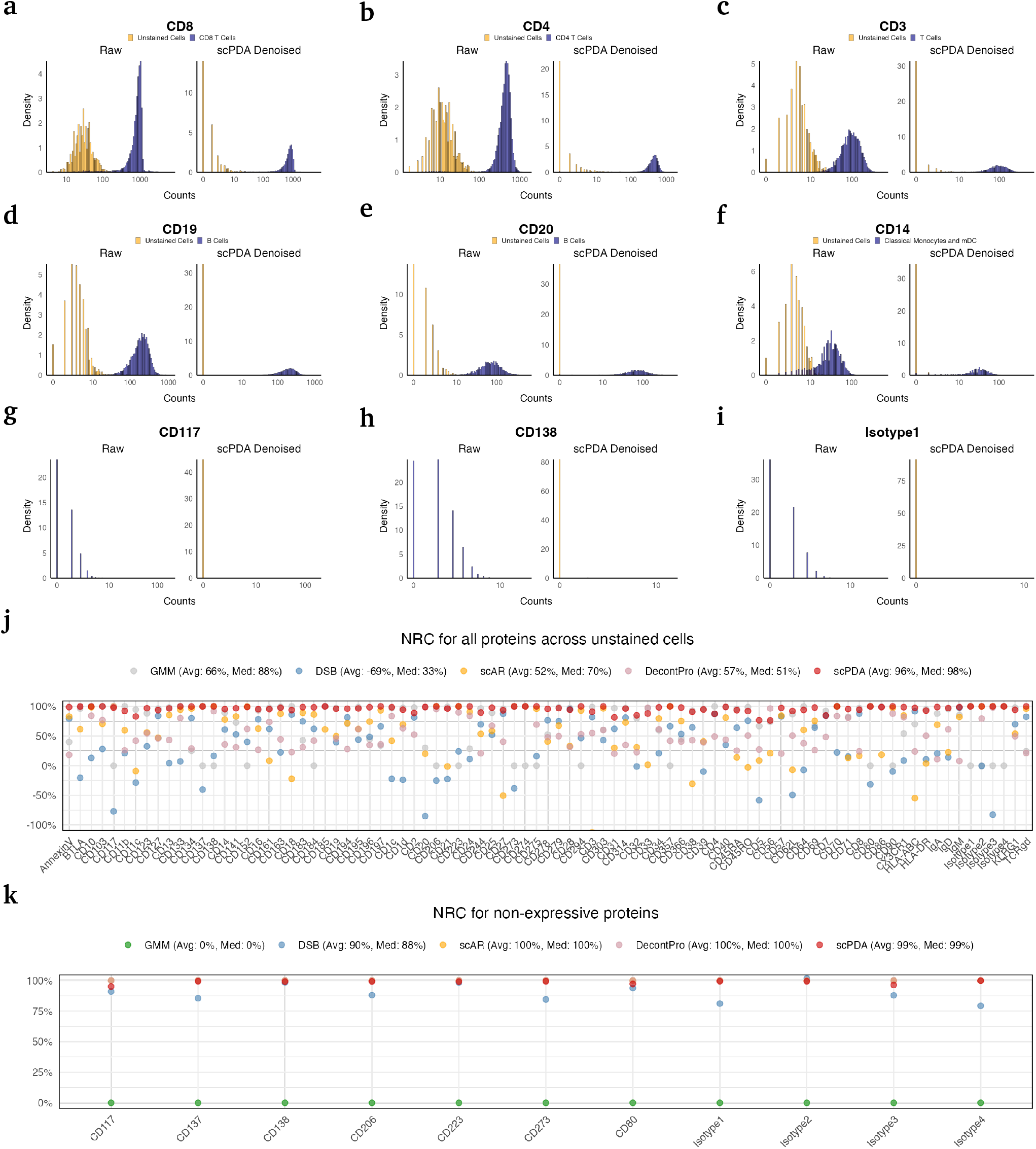
scPDA reduces the protein counts in unstained cells. **a-f**, The density plots displaying the counts of CD8(**a**), CD4(**b**), CD3(**c**), CD19(**d**), CD20(**e**), and CD14(**f**) within unstained cells versus corresponding expressive cell population, before (Left) and after (Right) scPDA denoising. **g-h**, The density plots displaying the counts of non-expressive proteins CD117(**a**), CD138(**b**), and Isotype1(**c**), before (Left) and after (Right) scPDA denoising. **j** The NRC on all 87 antibodies for unstained cells, derived from GMM, DSB, scAR, DecontPro, and scPDA. The average (Avg) and median (Med) NRC of each method is annotated. **k** Similar to **(j)**, the NRC is achieved on non-expressive proteins.

scPCA dramatically reduces the noise level, as shown in the right panels of Figures 1a-i. Considering CD8 again, its average counts in unstained cells decreased to 4.22, which represents an 88% decrease. At the same time, its average count in its targeted cell type (CD8 T cells) is barely changed. This indicates scPCA’s ability to accurately distinguish noise from the signal and to reduce the noise without noticeably shrinking the signal.

To quantitatively measure the performance of denoising, we borrow the concept from materials science and define the noise reduction coefficient (NRC) for a protein as

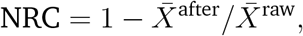

where 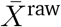 and 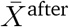 denote the average noise levels before and after applying the denoising method. Roughly speaking, the NRC represents the proportion of noise that has been eliminated. The ideal value of NRC is 1, and a larger NRC indicates a more thorough removal of noise. Typically, NRC is positive; however, if a denoising method amplifies rather than eliminates noise, the NRC may become negative, indicating the method’s poor performance.

Figures 1j show the NRC values for all proteins in the unstained cells. Results from our method, scPCA, as well as four other methods—GMM, DSB, scAR and DecontPro—are shown. Under the title of the figure, the mean and median NRC values from different methods are also provided. The mean and median values of our scPDA method are 96% and 98%, respectively, while the best other method is GMM, which yields mean and median values of 66% and 88%, significantly lower than our method. DSB performs the worst and even results in a negative mean value. The margin of performance difference between scPDA and other methods is remarkable in the unstained cells.

In the case of non-expressive proteins, as shown in Figure 1k, scAR, DecontPro, and our method, scPDA, all yield mean NRC values of 1 or close to 1, indicating perfect or near-perfect performance. Another method, DSB, also performs efficiently, with a mean NRC of 90%. However, GMM, which was the best performer among other methods for unstained cells, fails completely in this scenario, with a mean NRC of 0%. Such failure highlights GMM’s difficulty in handling uni-modal distributions, which are typical for non-expressive proteins; in such cases, GMM does not remove the only mode, which should be zero but appears at a non-zero location due to noise.

Overall, scPDA provides the most consistent and robust performance, and the lead of scPDA over other methods is pronounced.

### scPDA improves normalized data

Many subsequent analyses of single-cell protein data rely on the Centered Log Ratio (CLR), which is a widely-used type of standardized data reflecting the relative expression level of a protein compared to other proteins in the same cell [3, 22]. We call such transformed data “CLR values,” and we have found that scPDA remarkably improves the quality of CLR values. Here, used as an example, is a CITE-seq dataset that profiled 30,672 bone marrow cells with a panel of 25 antibodies [9].

The most important use of single-cell surface-protein data is to classify cells into different known cell types, which is often done by checking the expression status of so-called “marker proteins.” Some proteins are expected to be expressed in certain cell type(s), in which case they are called “positive markers” for these cell types. They do not express in other cell types, in which case they can be referred to as “negative markers” or “non-markers.” Therefore, within a cell type, positive markers are expected to be expressed much more than negative markers. Unfortunately, this may not be the case in real data due to the presence of noise.

We can easily identify such examples in Figures 2a-f. Each figure illustrates the distribution of CLR values for all 25 proteins in one of the six cell types. For instance, CD161, a C-type lectin-like receptor, is known as a positive marker for NK cells, while CD3 is known as a negative marker for these cells. However, the mean CLR value of CD161 is lower than that of CD3 (0.90 vs 0.93) in NK cells, as shown in Figure 2a, where the protein names CD161 and CD3 are highlighted in red for easy identification.

**Figure 2:**
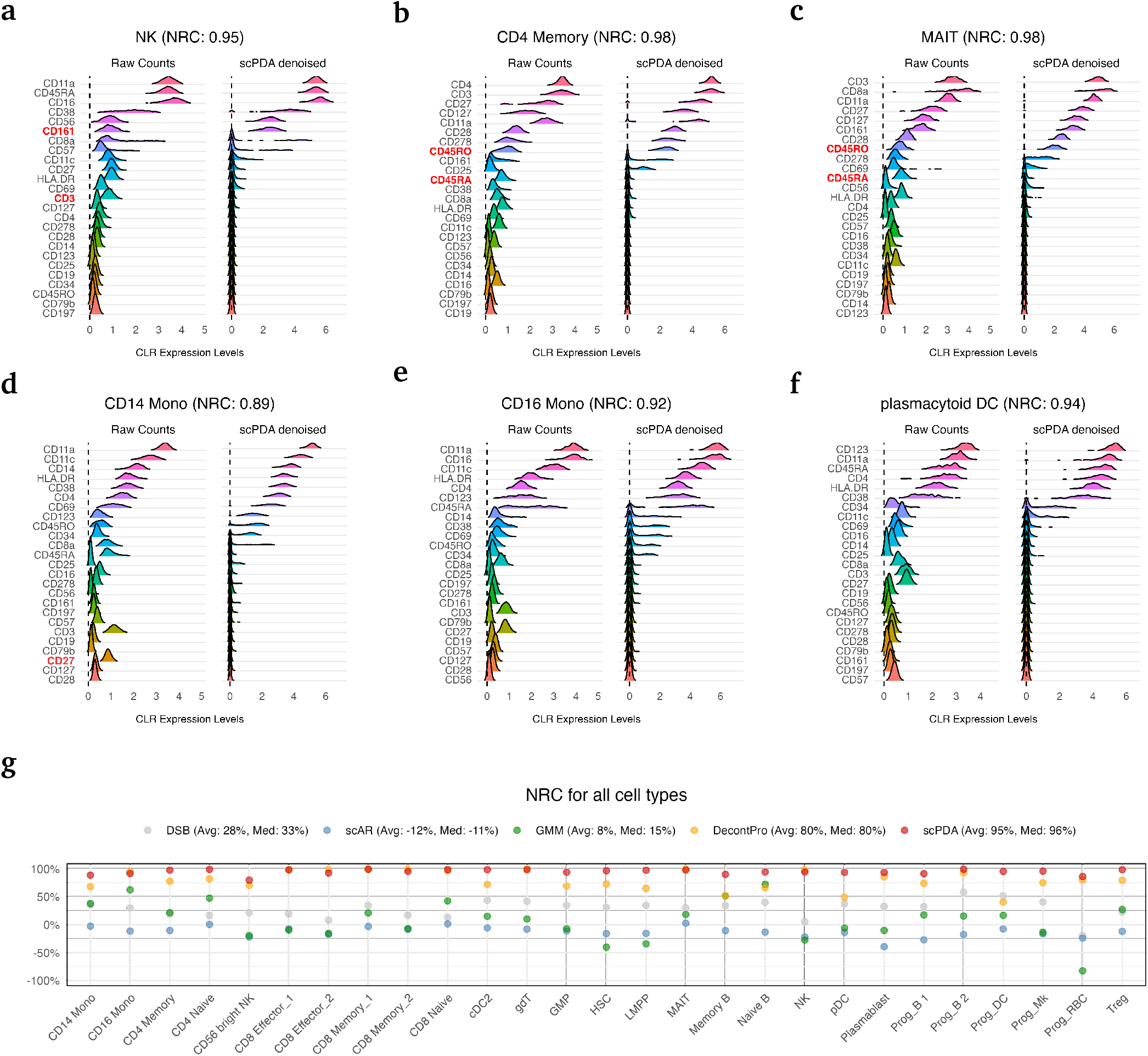
scPDA’s efficacy in noise correction and compatibility with normalization. **a-f**, Ridge plots displaying the CLR values of each protein within NK cells **(a)**, CD4 Memory T Cells **(b)**, MAIT cells **(c)**, CD14 Monocytes **(d)**, CD16 Monocytes **(e)**, and plasmacytoid dendritic cells (pDC) **(f)**, respectively, derived from raw counts (Left) vs scPDA-denoised counts (Right). The corresponding NRC is annotated at the top. **g**, The NRC on all cell types, derived from DSB, scAR, GMM, DecontPro, and scPDA. The average (Avg) and median (Med) NRC of each method is annotated.

As another example (shown in Figure 2b), CD45RO and CD45RA (protein names highlighted in red in the figure) are known as the only markers in the 25-antibody panel that differentiate Naive T cells and Memory T cells. The former is expected to be highly expressed in Memory T cells, while the latter is expected to be highly expressed in Naive T cells. However, in real data, due to noise, the expression of the former is not always significantly higher than that of the latter in Memory T cells. Specifically, the mean CLR value of CD45RO is only marginally higher than that of CD45RA in a type of Memory T cell (CD4 Memory cells), being 0.95 and 0.79, respectively. What is even more problematic (shown in Figure 2c), the mean CLR value of CD45RO is even lower than that of CD45RA in another type of Memory T cell (MAIT cells, which are a subset of CD8 Memory T cells), being 0.79 and 0.92, respectively.

scPDA remarkably improves the CLR values. It is able to differentiate between noise and signal, reducing the CLR values of noise while enlarging the CLR values of the signal, resulting in a much clearer separation between the two. In NK cells (Figure 2a), the mean CLR value of CD3 is reduced from 0.93 to 0.12, while that of CD161 is increased from 0.90 to 2.24. The mean CLR value of CD45RA in CD4 Memory cells (Figure 2b) is reduced from 0.79 to 0.14, and in MAIT cells (Figure 2c) from 0.92 to 0.24, while the mean CLR value of CD45RO in CD4 Memory cells (Figure 2b) is increased from 0.95 to 2.24, and in MAIT cells (Figure 2c) from 0.79 to 1.90. As an additional example, CD27, known as a pan-T-cell marker, exhibits moderate CLR values within non-T cells prior to denoising (e.g., 0.89 in CD14 Monocytes, Figure 2d); after scPDA denoising, its value is reduced to nearly zero (e.g., 0.03 in CD14 Monocytes, Figure 2d).

To quantitatively assess the denoising efficiency of a method, we define the NRC for a cell type as the average NRC of all its negative marker proteins. As before, a larger NRC value indicates more thorough denoising, with a value of 1 meaning perfect denoising.

The NRC values for different cell types using our method, scPDA, as well as four other methods—GMM, DSB, DecontPro, and scAR—are shown in Figure 2g. Three of the other four methods perform poorly. scAR has negative NRC values in most cell types, with an average of −12%, indicating that it falsely increases the noise of negative markers. GMM and DSB show moderate denoising effectiveness, achieving average NRC values of 8% and 28%, respectively. Among the other four methods, DecontPro provides much better performance, achieving an average NRC value of 80%. On the other hand, our method, scPDA, consistently yields remarkably higher NRC values, ranging from 80% to 99%, with an average of 95%. This indicates that scPDA is much more effective in removing noise than the others.

### scPDA enhances the gating strategy

Cell type annotation based on surface protein expression, a major use of such data, is often accomplished using the gating strategy. This strategy is simple and straightforward; it determines a cell’s cell-type identity by checking the expression of one or a few marker genes. For example, CD19 is a known positive marker for B cells. If its expression in a cell exceeds a pre-set threshold, then the cell is labeled as a B cell; otherwise, the cell remains unlabeled. However, the presence of noise may reduce the efficiency of the gating strategy by blurring the separation between different cell types based on their expression of marker genes.

scPDA may enhance the efficiency of the gating strategy by sharpening the separation in expression. To demonstrate this, we used a CITE-seq dataset [21] that profiles PBMCs with a 188-antibody panel. The author of the data employed a gating strategy based on the expression of CD3, CD19, CD4, CD8, CD14, CD16, and CD56 to categorize PBMCs into five major cell types: CD4-T cells, CD8-T cells, B cells, Natural Killer (NK) cells, and Classical Monocytes (CM). Detailed criteria for cell type definition using the gating strategy are provided in Supplementary Material section S4.

The top left panel of Figure 3a shows an example of blurred expression before denoising. It is a bi-axial scatter plot depicting the CLR values of CD4 against CD14. CD4 is expected to be highly expressed in CD4-T cells, expressed at a low level in CM cells, and not expressed in other cell types. As shown in the figure, although CD4 exhibits clearly higher expression in CD4-T cells, its expression in CM cells shows a moderate overlap with other cells (i.e., neither CD4-T nor CM). There is no threshold in the CD4 expression that effectively separates CM cells from others. The problem with CD14 expression is even more severe. CD14 is expected to be expressed in CM but not in other cell types. However, a significant group (455 cells) of CD4-T cells shows moderately high CD14 expression. The average expression of these 455 CD4-T cells is 0.32, not too far from the average CM expression, which is 0.58. In this case, there is no satisfactory threshold for gating using CD14, as a low threshold will mislabel some CD4-T cells as CM, while a high threshold will leave some CM cells unclassified.

**Figure 3:**
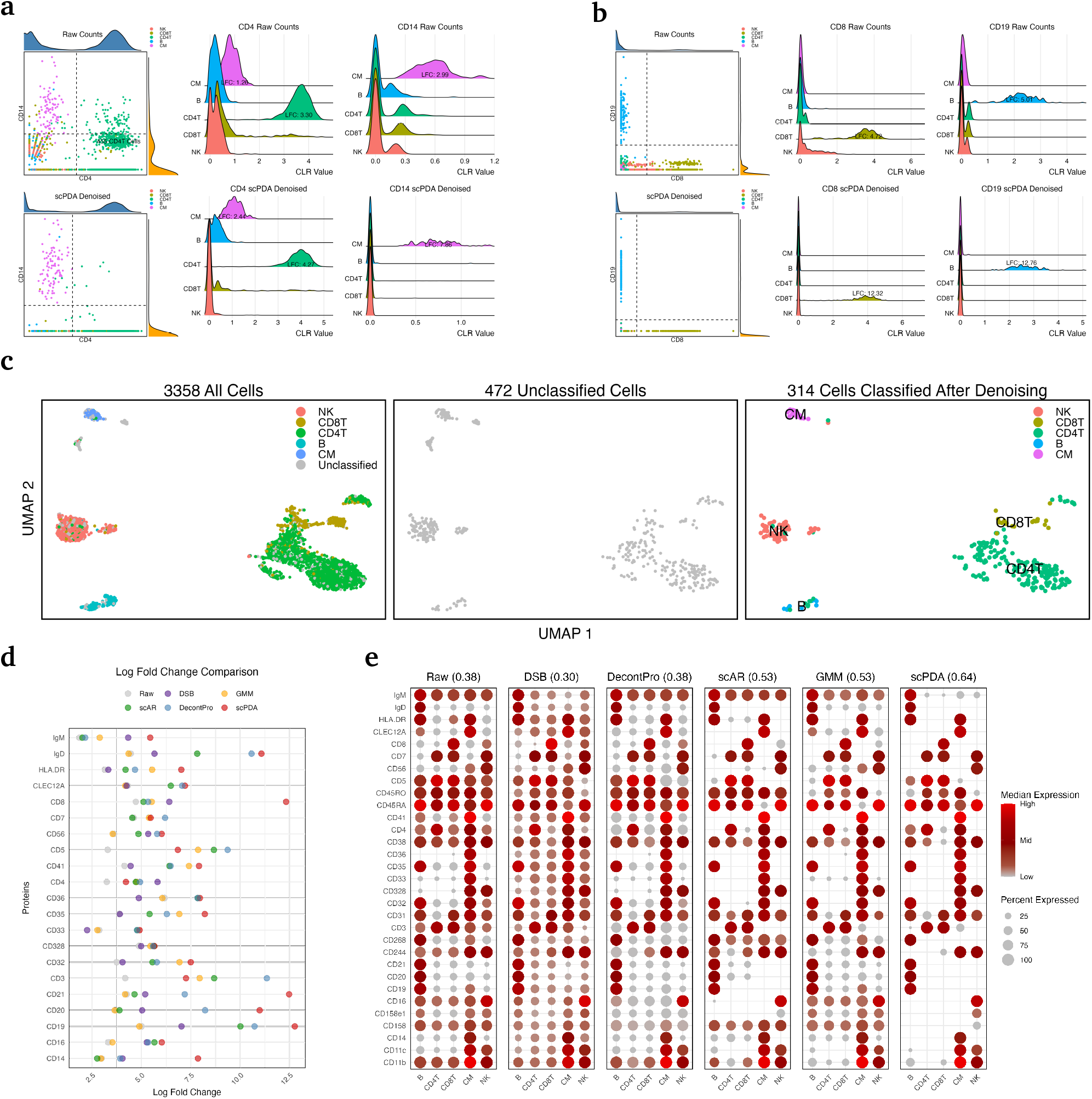
scPDA improves protein-based gating strategy. **a**, (Left) The bi-axial plot of the expression of CD4 and CD14, derived from raw counts (Top) and scPDA-denoised counts (Bottom), with colors correspond to the cell types. (Middle) The ridge plots of the expression of CD4 derived from raw counts (Top) and scPDA-denoised counts (Bottom), with the Log2 Fold Change (LFC) annotated. (Right) Similar to (Middle), the ridge plots is based on the expression of CD14. **b**, Similar to **a**, but compares the expression level of CD8 and CD19. **c**, (Left) UMAP visualization of classified cells on the gene space. Cells are colored by the annotated cell types. (Middle) Similar to (left), but displaying unclassified cells that failed the gating strategy. (Right) Previously unclassified cells but now pass the gating strategy after scPDA denoising, colored by the assigned cell type annotation. **d**, LFCs of each protein, derived from raw counts, DSB, GMM, scAR, DecontPro, and scPDA, respectively. **e**, Dotplots by Raw, DSB, DecontPro, scAR, GMM, and scPDA, respectively. The dot color displays median expression per protein (column) within each cell type (row), and the dot size represents the percentage of cells with expression level greater than zero. The corresponding sparsity score of each method is annotated.

The top left panel of Figure 3b shows another such example. CD8 is a (positive) marker for CD8-T cells, but there is a moderate overlap between its expression in CD8-T and NK cells, which is clearly illustrated in the top middle panel of Figure 3b.

scPDA makes the gating strategy much easier and more efficient. In the example shown in Figure 3a, the CD14 expression of the 455 CD4-T cells that previously showed moderately high expression is reduced to nearly zero. More specifically, only 11 of them exhibit a CD14 expression greater than 0.2, making the distinction between CM and CD4-T cells using the gating strategy on CD14 straightforward and less error-prone. In the example shown in Figure 3b, 64 NK cells had CD8 expression greater than 10 before denoising. After denoising, only one retains an expression level greater than 10.

Such examples are not rare. Overall, prior to denoising, 472 out of 3,358 cells (14.1%) were reported as unclassified by the author using the gating strategy. After denoising with scPDA, 314 of these 472 unclassified cells (66.5%) passed the same gating strategy with the same thresholds and received cell-type labels. We wish to assess the accuracy of the cell-type labels of these newly assigned cells, although the ground-truth labels are unknown. We propose using cell clustering derived from the RNA modality for an independent and unbiased assessment. The results are illustrated in the three panels of Figure 3c. In the left panel, the cells are positioned in a two-dimensional space given by applying UMAP to the RNA modality, and they are colored according to their cell-type labels derived from applying the gating strategy to the original protein modality without denoising (these labels are provided in the paper that generated the data). We see that cells assigned to the same cell type largely cluster together. The 472 cells unclassified by the gating strategy in the original data are shown independently in the middle panel. Among these 472 cells, the 314 cells that pass the gating strategy in the scPDA denoised data are depicted in the right panel, colored according to their newly assigned cell types by the gating strategy. It is clear that in the right panel, cells of the same color still cluster together, and the colors align closely with those in the left panel. This indicates that most of these newly assigned labels are correct.

### scPDA enhances the differential expression analysis

scPDA enhances the power of differential expression analysis of protein expression between different cell groups by amplifying the Log2 Fold Change (LFC). We illustrate this using the same PBMC dataset as in the previous section.

Still consider the distribution of CD4 expression, which is expected to be trimodal: high in CD4-T cells, moderate in CM, and low in other cell types. The middle panel of Figure 3a shows the distribution of CD4 expression before (top) and after (bottom) denoising using scPDA. Note that the y-axis (density) of the top plot and bottom plot are on different scales, since the expression of unexpressed proteins gets much closer to zero after denoising, causing their density around zero to increase significantly. Due to this scale difference, the peak corresponding to proteins that should be highly expressed (e.g., the green peak in the middle panel) appears much lower in the denoised data than in the original data, although the density it represents remains roughly unchanged after denoising.

As shown in the middle panel of Figure 3a, the LFC of CD4 expression increases from 3.30 to 4.27 in CD4-T cells compared to the unexpressed cell types, and from 1.26 to 2.44 in CM cells. The right panel of Figure 3a, the middle panel of Figure 3b, and the right panel of Figure 3b provide three more examples, where the LFC increases from 2.99 to 7.86, 4.72 to 12.32, and 5.01 to 12.76, respectively. Note that these fold changes are in the Log2 scale, and thus these increases are quite dramatic. For example, a change of LFC from 4.72 to 12.32 means a 194-fold change on the original scale.

Figure 3d provides a comprehensive comparison of the LFC derived from all marker proteins using different methods: raw data, DSB, GMM, scAR, DecontPro, and scPDA. Compared to the raw data, which has an average LFC of 3.92, DSB, GMM, scAR, DecontPro, and scPDA have average LFCs that are 0.87, 1.16, 1.77, 2.72, and 3.94 larger, respectively. Again, scPDA outperforms other methods substantially.

As a further comparison of scPDA’s contribution to differential expression analysis relative to other methods, we construct a dot plot for each method, as illustrated in Figure 3e. In these dot plots, the color of each dot represents the median protein expression within a cell type, while the size of the dot indicates the proportion of cells within that cell type exhibiting an expression level no less than 1. Note that we use no less than 1 instead of larger than 0 since some methods never output exact zeros for non-zero counts.

Scrutinizing these plots, we find that DSB appears to blur the distinction between noise and signal in some cases, with increased detection of proteins in unrelated cell types, such as CD19 and CD20 (B cell markers) in non-B cells. In such cases, DecontPro does not improve much, as indicated by the similar noise level in non-marker proteins. scAR and GMM, on the other hand, effectively removes the noise in these cases, as evident from the decreased CD8 expression in NK cells when compared to raw, DSB, and DecontPro. scPDA further outperforms scAR and GMM, as demonstrated by the reduced expression of the B-cell marker IgM in non-B cells and the NK-cell marker CD16 in non-NK cells.

To summarize each heatmap, we define a sparsity score as the percentage of dots with expression below 1. Since proteins are expected to have zero expression in all cell types where they are not markers, a higher sparsity score indicates that more non-markers have zero expression, making the markers stand out, which is favored. Therefore, this sparsity score provides a quantitative measure of the thoroughness of denoising. scPDA achieves a significantly higher sparsity score (0.64) compared to other methods, which have sparsity scores of 0.53 or lower.

### scPDA refines the estimation of background probability

When determining cell types, the gating strategy is typically conducted manually. For a marker protein, this strategy involves manually choosing a threshold; if the expression is higher than the threshold, it is regarded as expressed, and the cell is assigned to the “positive population.” Conversely, if the expression is lower than the threshold, it is regarded as unexpressed, and the cell is assigned to the “negative population.” This manual strategy introduces arbitrariness due to the choice of the threshold. Methods such as GMM and our method, scPDA, potentially provide an automatic alternative, as they offer an (estimated) background probability, which is the probability of an expression coming from background noise rather than a biological signal.

We can use the AUC (area under the ROC curve) to assess the quality of the estimated background probability by different methods, particularly by GMM and scPDA. The other three methods, DSB, DecontPro, and scAR, do not provide an estimate of the background probability. To still include them in the comparison, we obtained their estimated background probabilities by fitting GMMs on the denoised counts given by these three methods.

In this section, we will demonstrate scPDA’s advantage in providing a better background probability using a dataset generated by the tri-omic technology TEA-seq [5]. This dataset measures three modalities of 18,011 PBMCs: gene expression, chromatin accessibility, and the abundance of 46 proteins. We will report the AUCs of different methods, as well as the LFCs.

Figure 4a provides four density plots of CD3 expression, where the positive population (T cells) and the negative population (other cells) are colored differently. These plots are derived from DSB, DecontPro, scAR, and scPDA, respectively. Gating on this protein to identify T cells is not a challenging task, as indicated by an AUC of 0.97 even using the raw data. DSB, DecontPro, and scAR returned similar AUCs of 0.97, 0.98, and 0.97, respectively, while scPDA further improves the AUC to 1.00. The LFC is 0.85 on raw counts and increases to 2.20, 1.08, 1.27, and 3.53 after denoising by DSB, DecontPro, scAR, and scPDA, respectively. scPDA still performs the best in this aspect. Figure 4b shows another protein, CD19. While all methods yield a perfect AUC (1.00), scPDA achieves a much larger LFC than the other methods.

**Figure 4:**
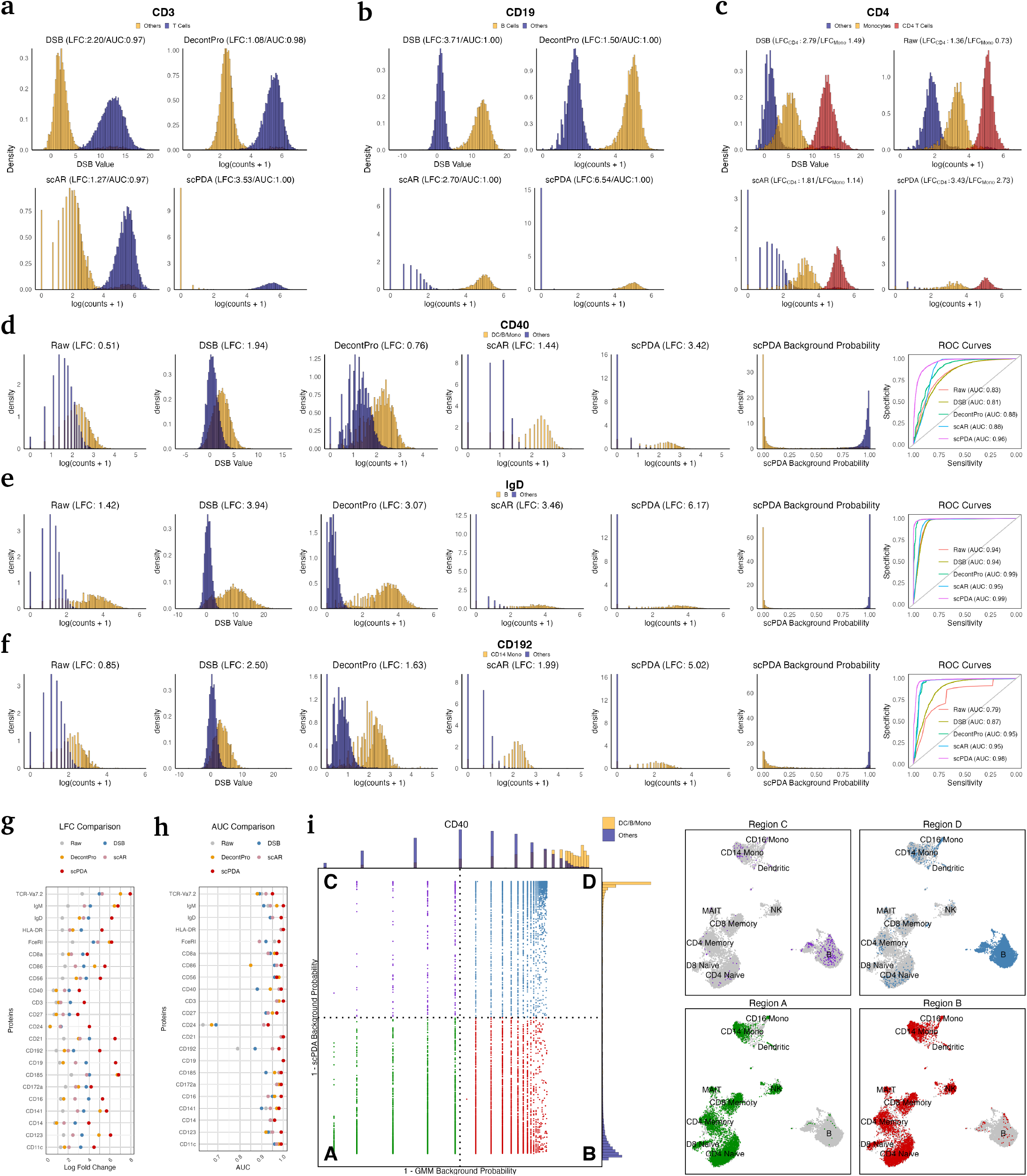
Enhanced background probability estimation with scPDA. **a**, Four density plots displaying CD3 expression within positive population (T cells) versus negative population (non-T cells), derived from DSB, DecontPro, scAR and scPDA. Log2 Fold Change (LFC) and area under the ROC curve (AUC) values are annotated. **b-c**, Analogous to **a**, but focusing on the expression of CD19 (**b**) and CD4 (**c**). **d-f**, Similar to **a**, but including three additional plots for CD40 (**d**), IgD (**e**), and CD192 (**f**): raw count density plot, scPDA background probability density plot, and ROC Curve comparisons among different methods. For DSB, DecontPro and scAR, ROC Curves are obtained based on the background probability derived from a two-component GMM fit on denoised counts. **g**, LFCs for each protein between its corresponding positive and negative populations, obtained from raw counts, DSB, DecontPro, scAR, and scPDA. **h**, Similar to **g**, but comparing AUC for each protein across denoising methods. **i**, (Left) Scatter plot of 1 - GMM estimated background probability (x-axis) against 1 - scPDA estimated background probability (y-axis), with corresponding density plots attached on top and right side. Dashed lines represent the 50% background probability threshold for GMM and scPDA, segmenting the plot into four quadrants. (Right) UMAP visualizations of cell populations using Seurat’s WNN mRNA-protein multimodal algorithm, highlighting cells from each quadrant in the scatter plot.

**Figure 5:**
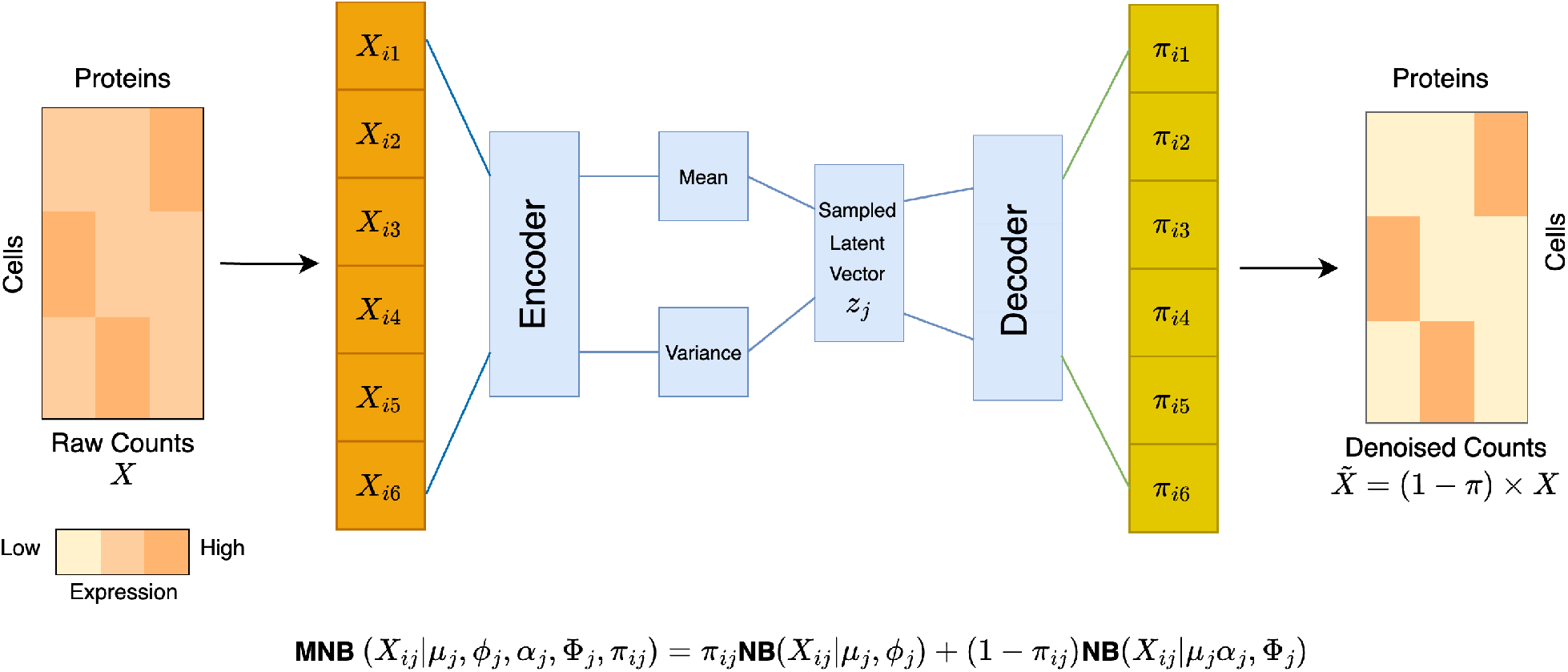
Architecture of scPDA. *X*_*ij*_, the observed count of protein *j* on cell *i*, is modeled using a two-component negative binomial mixture distribution, containing parameters *π*_*ij*_, the mixture weight, (*µ*_*j*_, *ϕ*_*j*_) and (*µ*_*j*_*α*_*j*_, Φ_*j*_), the mean and dispersion for component of background noise and biological signal, respectively. The protein count vector for each cell *i* is converted by an encoder neural network to the parameters of the distribution that latent variable 𝒵_*j*_ follows. The decoder neural network then transforms sampled 𝒵_*j*_ to *π*_*j*_. The denoised count matrix, 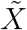, is given by (1 − *π*)*X*.

Figure 4c displays protein CD4, which is expected to exhibit three peaks: high in CD4 T cells, moderate in monocytes, and zero in others. In this three-class case, the AUC is not applicable, and therefore, we report only the LFCs (LFC of CD4 T versus others, and LFC of monocytes versus others). Similar to the observations in Figure 3a, scPDA, despite assuming a two-component-mixture model, effectively handles data with more than two modes. After denoising by scPDA, the tri-modal distribution is not only well preserved but also becomes significantly clearer. This is evident as the LFCs produced by scPDA are substantially larger than those of the raw data and the denoised data from other methods.

Figure 4d illustrates a much more challenging case: CD40, whose expression shows significant overlap between the positive population (B cells, dendritic cells (DC), and monocytes (Mono)) and the negative population (other cells). In the raw data, the AUC is 0.83. DSB fails to improve the AUC, yielding an AUC of 0.81. scAR and DecontPro moderately elevate the AUC to 0.88. In contrast, our method, scPDA, increases the AUC to 0.96, marking a substantial improvement. Figure 4d also presents the LFCs of different methods, with scPDA significantly outperforming the others in this aspect.

In Figure 4d, the distributions of the background probability for CD40, estimated by scPDA, are shown, colored to indicate whether they are DC/B/Mono cells or not. The vast majority of DC/B/Mono cells have background probabilities below 0.1, while the vast majority of other cells have background probabilities above 0.9. This demonstrates that scPDA not only accurately distinguishes between the two groups of cells but also does so with high confidence.

Similar situations occur with other proteins that are not as dramatically affected but are still significant, such as IgD, a B cell marker, and CD192, a marker for CD14 Monocytes, as illustrated in Figures 4e-f. The LFCs and AUCs for all 22 marker proteins from various methods are shown in Figures 4g-h. The average LFCs provided by raw data, DSB, DecontPro, scAR, and scPDA are 1.42, 3.34, 3.05, 2.68, and 5.21, respectively; similarly, the average AUCs are 0.93, 0.93, 0.95, 0.95, and 0.98, respectively. Again, our method, scPDA, demonstrates a substantial improvement in performance compared to other methods under both evaluation criteria.

Using CD40 as an example, we present a closer examination to show how scPDA better identifies the negative and positive populations than a two-component GMM. On the left side of Figure 4i, a scatter plot depicts the 1 minus background probability estimated by scPDA versus that estimated by GMM. The horizontal and vertical black dotted lines correspond to the 50% background probability threshold derived from scPDA and GMM, respectively. These two lines divide the plot into four regions, which are marked in the figure: (region A) cells classified as negative (i.e., in the negative population of CD40) by both GMM and scPDA, (region B): cells classified as positive by GMM but as negative by scPDA, (region C): cells classified as negative by GMM but as positive by scPDA, and (region D): cells classified as positive by both GMM and scPDA. So we see that scPDA and GMM agree in regions A and D but disagree in regions B, which contains 5,655 cells (32% of all cells), and region C, which contains 289 cells (2% of all cells).

To determine which method’s results are likely correct in regions B and C, we borrow information from the RNA modality of the data. Combining the RNA modality and the protein modality using Seurat’s weighted nearest neighbor algorithm [9], the cells are positioned in a two-dimensional UMAP space in the right panel of Figure 4i. In the four subfigures of the right panel of Figure 4i, the cells (i.e., the points) are the same, but in each subfigure, cells in one region (A, B, C, or D) are colored in the same color as they are in the left panel of Figure 4i. According to the left panel of Figure 4i, GMM indicates that the purple (region C) and green (region A) cells should be of the same cell type(s), while the blue (region D) and red (region B) cells should be of the same cell type(s). From the right panel of Figure 4i, this is very unlikely the case. In contrast, scPDA indicates that the purple (region C) and blue (region D) cells are of the same cell type(s), while the green (region A) and red (region B) cells are of the same cell type(s). From the right panel of Figure 4i, this appears to be the case. This result strongly suggests that scPDA’s estimations of background probability are more accurate than those of GMM. Given that regions B and C, where the two methods diverge, comprise as much as 34% of all cells, the superiority of scPDA is substantial.

## Discussion

The expression of surface proteins provides an easy way to identify cell types, yet widespread noise deteriorates the performance of the gating strategy. Recent denoising methods suffer from various limitations. In this study, we introduced scPDA, a novel probabilistic model that overcomes these limitations. We applied scPDA to multiple single-cell protein datasets and demonstrated its remarkable improvement in noise reduction, normalization, gating strategy, differential expression analysis, and background probability estimation.

scPDA is light in computation and memory use, making it scale well to datasets with large numbers of cells. For instance, in the PBMC dataset used in the first subsection of the Results section, which contains over 50,000 cells and 87 proteins, scPDA completed the computation in about 10 minutes and required 530MB of RAM. In comparison, scAR also required about 10 minutes and consumed 552MB of RAM. DSB completed its processing in about 1 minute with 367MB of RAM. Conversely, DecontPro consumed significantly more resources, requiring over 38 hours of computational time and 360GB of RAM. Detailed results can be found in Supplementary Material Figure S5.

scPDA assumes a two-component mixture model, which can raise concerns when the protein counts display either a single mode or more than two modes. Single-mode cases typically occur with non-expressive proteins. As described in the Methods section, scPDA incorporates a penalty term in its target function to enhance the distinction between the centers of the two peaks. In practice, scPDA usually assigns the single peak as the background and removes it, effectively dealing with non-expressive proteins. However, scPDA may be less suitable for a protein expressed across the entire cell population, although such cases are very rare. Among all the datasets we have analyzed, only one contains such instances (antibodies CD18 and HLA-ABC in the dataset [20]). Instances of more than two modes also occur in real data, as seen in Figures 4c and 3a. In these cases, scPDA typically successfully identifies the mode corresponding to the background noise as the lower mode and removes it.

## Methods

### The scPDA model

#### Variational Autoencoders (VAEs)

VAEs [23, 24] evolve from traditional autoencoders [25], which are artificial neural networks designed for unsupervised learning, compressing and reconstructing data through an encoder and a decoder. Traditional autoencoders minimize reconstruction loss, typically measured by mean squared error, but they suffer from discontinuities in the latent space, leading to potential decoding errors [26]. VAEs [23, 24] address this issue by enforcing a probabilistic distribution in the latent space. This strategy results in a structured and continuous latent space, allowing for more meaningful and controlled data generation. In addition to reconstruction loss, VAE also minimizes the Kullback-Leibler (KL) divergence between the distribution of latent representation and a prior distribution, which is usually a standard Gaussian 𝒩 (0, 𝕀).

#### The probabilistic model of scPDA

scPDA models the observed count *X*_*ij*_ of protein *j* in cell *i* by a two-component negative binomial mixture (NBM) distribution

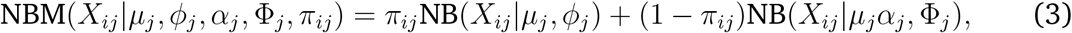

where NB(*x*|*µ, ϕ*) denotes the probability function of a negative binomial distribution with mean *µ* and dispersion *ϕ*, i.e.,

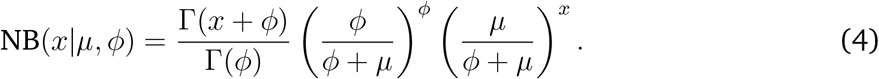

In equation 3, *π*_*ij*_ is the background probability, representing the likelihood of *X*_*ij*_ originating from noise. scPDA uses two distinct negative binomial distributions, with means *µ*_*j*_ and *α*_*j*_*µ*_*j*_ and dispersions *ϕ*_*j*_ and Φ_*j*_, to model the background noise and biological signal, respectively. *α*_*j*_ is expected to be larger than 1, which will be reinforced using an activation function. Notably, the mean and dispersion parameters in scPDA are protein-specific and not cell-specific.

#### The VAE architecture of scPDA

The parameters of the NBM model are determined in three distinct fashions. First, scPDA employs the following VAE framework to infer *π*_*ij*_:

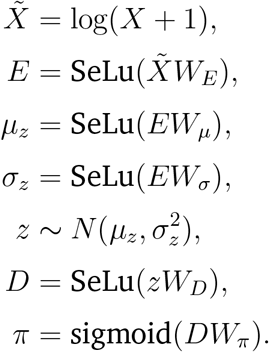

Here, *X* is the observed count matrix, *E* and *D* represent the encoder and decoder layers, and 𝒵 is the latent variable that follows a normal distribution parameterized by 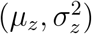.

Subsequently, the protein-specific parameters (*ϕ*_*j*_, *α*_*j*_, Φ_*j*_) are registered as the global network parameters and updated through backward propagation during optimization. To ensure the positivity of *ϕ*_*j*_ and Φ_*j*_, and to ensure that *α*_*j*_ *>* 1, activation functions are chosen as:

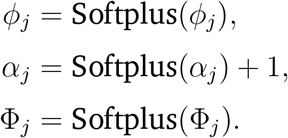

Finally, *µ*_*j*_, the expectation of the background noise, is estimated before the training of VAE. It is set as the lowest peak of a GMM fitted on the raw counts of protein *j*. Note that the GMM is not fitted on the log-transformed data, in order to avoid the problem of artificial peaks in the low end caused by logarithmic transformation. Also, both GMM models with two and three components are fitted on the data, and the one with a lower BIC is selected. This strategy for estimating *µ*_*j*_ helps narrow the optimization search space and prevents entrapment in local minima of poor quality when training VAE.

#### The loss function of scPDA

scPDA’s loss function comprises three components

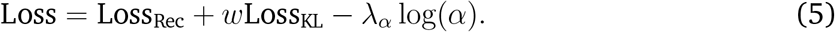

The first component is the reconstruction loss

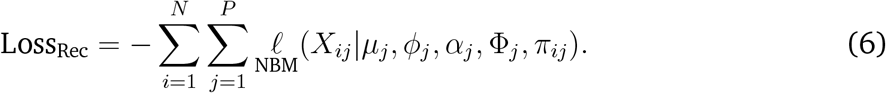

This loss measured by the negative log likelihood of the NBM distribution, representing how closely the decoder’s output matches the original input.

The second component is the KL Divergence

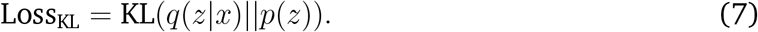

The KL Divergence quantifies the deviation of the learned latent representation *q*(𝒵|*x*) from the prior distribution *p*(𝒵), which is a standard Gaussian 𝒩 (0, 𝕀). The inclusion of this loss serves as a crucial regularization in the latent space, ensuring that similar latent points yield similar outputs.

As the third component, scDPA introduces a penalty term proportional to log(*α*). This is motivated by the cases where a protein is not expressed on whole cell population, which result in a uni-modal distribution that only contains background noise. The penalty term can address such problem by enforcing the potential biological signal significantly exceed background noise.

These three components are combined using weights 𝓌 and *λ*_*α*_.

#### Inference

The output of scPDA, *π*_*ij*_, is interpreted as the probability of protein *j* not expressed on cell *i*, and 1 − *π*_*ij*_ as the probability of it being expressed. One can call protein *i* to be present in cell *j* if 1 − *π* is over a threshold and absent otherwise. In scPDA, the denoised data is given by

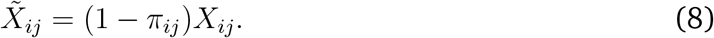

#### Training and hyperparameter selection

scPDA is trained using the Adam optimizer [27], with the learning rate scheduled to exponentially decay by 0.99 every epoch. The total number of training epochs is set to 500. To determine a default set of other hyperparameters for scPDA, we conducted a grid search on three single-cell protein expression datasets [9, 21, 5]. The hyperparameters considered include number of nodes in each layer, batch size, initial learning rate, 𝓌, and *λ*_*α*_. For each dataset, we evaluated their combinations and selected the one yielding the highest average LFC across all datasets (Supplementary Material Figure S4) as the default setting of hyperparameters, which are: 100 units in the first layer, 50 units in the second layer, 15 units in the latent space, a batch size of 256, an initial learning rate of 0.005, 𝓌 set to 0.25, and *λ*_*α*_ at 0.1. The result of all data analysis presented in our paper were derived from these default parameters.

#### Definition of empty droplets

In [16], two types of empty droplets were provided by the author of DSB, defined by cell hashing and library size threshold, respectively. We chose the former as it provides a better noise estimation for DSB, scAR and DecontPro. In [5], the definition of empty droplets is also given by the author of DSB, based on the library size threshold on both the protein and gene modalities.

#### Benchmark against GMM

GMM is fit on the log counts independently for each protein using the R package `mclust` [28]. As suggested by the study of DSB, the number of components is chosen between two and three based on BIC. The denoised data of GMM is given by Equation 8.

#### Benchmark against DSB

DSB is implemented using the R package dsb, which offers two distinct implementations tailored respectively to scenarios with and without empty droplets. For cases without empty droplets, we utilized `ModelNegativeADTnorm`(), while `DSBNormalizeProtein`() was employed when empty droplets were available. Both functions include the parameter `use.isotype.control`, which is set to TRUE if isotype control antibodies are available, and FALSE otherwise. The output of DSB are normalized data, with the values representing the number of standard deviation above the noise level in empty droplets. These values can often be negative. Following the DecontPro paper [18], we exponentiated them so that they are comparable to denoised expression values given by other methods, which are non-negative.

#### Benchmark against scAR

scAR is implemented using the Python package scar, which also has two versions of implementation for the case with and without empty droplets. When empty droplets are available, the parameter `ambient_profile` is set to the proportion of each protein’s average expression in empty droplets. Otherwise, it is set to `None`, and scAR averages all the cell-containing droplets instead of empty droplets, to achieve the ambient frequency *α* in equation 1. The other parameters were set to their default values.

#### Benchmark against DecontPro

DecontPro, implemented using the R package decontX, requires cell cluster labels as inputs. To obtain these labels, we adhered to the workflow suggested by the author of DecontPro, utilizing the R Package Seurat [22] version 5.0.1. This involved the following steps: data normalization with `NormalizeData(normalization.method = “CLR”, margin = 2)`, dimension reduction using RunPCA`(npcs=10)`, neighborhood finding with FindNeighbors`(dims = 1:10)`, and cluster identification via `FindClusters()`. In our analysis, all parameters of DecontPro were kept at their default values.

## Data availability

The raw and preprocessed data used in [20, 5] can be downloaded at https://doi.org/10.35092/yhjc.13370915. The raw and preprocessed data in [9] can be accessed in the R package `SeuratData` [3] by calling `LoadData(ds = “bmcite”)`. The raw data in [21] are available in the National Center for Biotechnology Information (NCBI) Gene Expression Omnibus (GEO) repository, [Accession Number: GSE213282].

## Code availability

The scPDA package is available at GitHub https://github.com/PancakeZoy/scPDA.

## References

[1] Xiannian Zhang, Tianqi Li, Feng Liu, Yaqi Chen, Jiacheng Yao, Zeyao Li, Yanyi Huang, and Jianbin Wang. Comparative analysis of droplet-based ultra-high-throughput single-cell rna-seq systems. Molecular cell, 73(1):130–142, 2019.

[2] Robert Salomon, Dominik Kaczorowski, Fatima Valdes-Mora, Robert E Nordon, Adrian Neild, Nona Farbehi, Nenad Bartonicek, and David Gallego-Ortega. Droplet-based single cell rnaseq tools: a practical guide. Lab on a Chip, 19(10):1706–1727, 2019.

[3] Marlon Stoeckius, Christoph Hafemeister, William Stephenson, Brian Houck-Loomis, Pratip K Chattopadhyay, Harold Swerdlow, Rahul Satija, and Peter Smibert. Simultaneous epitope and transcriptome measurement in single cells. Nature methods, 14(9):865–868, 2017.

[4] Eleni P Mimitou, Caleb A Lareau, Kelvin Y Chen, Andre L Zorzetto-Fernandes, Yuhan Hao, Yusuke Takeshima, Wendy Luo, Tse-Shun Huang, Bertrand Z Yeung, Efthymia Papalexi, et al. Scalable, multimodal profiling of chromatin accessibility, gene expression and protein levels in single cells. Nature biotechnology, 39(10):1246–1258, 2021.

[5] Elliott Swanson, Cara Lord, Julian Reading, Alexander T Heubeck, Palak C Genge, Zachary Thomson, Morgan DA Weiss, Xiao-jun Li, Adam K Savage, Richard R Green, et al. Simultaneous trimodal single-cell measurement of transcripts, epitopes, and chromatin accessibility using tea-seq. Elife, 10:e63632, 2021.

[6] Di Ran, Shanshan Zhang, Nicholas Lytal, and Lingling An. scdoc: correcting drop-out events in single-cell rna-seq data. Bioinformatics, 36(15):4233–4239, 2020.

[7] Chenyang Xu, Lei Cai, and Jingyang Gao. An efficient scrna-seq dropout imputation method using graph attention network. BMC bioinformatics, 22:1–18, 2021.

[8] Geng Chen, Baitang Ning, and Tieliu Shi. Single-cell rna-seq technologies and related computational data analysis. Frontiers in genetics, 10:317, 2019.

[9] Tim Stuart, Andrew Butler, Paul Hoffman, Christoph Hafemeister, Efthymia Papalexi, William M Mauck, Yuhan Hao, Marlon Stoeckius, Peter Smibert, and Rahul Satija. Comprehensive integration of single-cell data. Cell, 177(7):1888–1902, 2019.

[10] Prabhu S Arunachalam, Florian Wimmers, Chris Ka Pun Mok, Ranawaka APM Perera, Madeleine Scott, Thomas Hagan, Natalia Sigal, Yupeng Feng, Laurel Bristow, Owen Tak-Yin Tsang, et al. Systems biological assessment of immunity to mild versus severe covid-19 infection in humans. Science, 369(6508):1210–1220, 2020.

[11] Jeffrey M Granja, Sandy Klemm, Lisa M McGinnis, Arwa S Kathiria, Anja Mezger, M Ryan Corces, Benjamin Parks, Eric Gars, Michaela Liedtke, Grace XY Zheng, et al. Single-cell multiomic analysis identifies regulatory programs in mixed-phenotype acute leukemia. Nature biotechnology, 37(12):1458–1465, 2019.

[12] Jin-Gyu Cheong, Arjun Ravishankar, Siddhartha Sharma, Christopher N Parkhurst, Simon A Grassmann, Claire K Wingert, Paoline Laurent, Sai Ma, Lucinda Paddock, Isabella C Miranda, et al. Epigenetic memory of coronavirus infection in innate immune cells and their progenitors. Cell, 186(18):3882–3902, 2023.

[13] Sergio Triana, Dominik Vonficht, Lea Jopp-Saile, Simon Raffel, Raphael Lutz, Daniel Leonce, Magdalena Antes, Pablo Hernández-Malmierca, Diana Ordoñez-Rueda, Beáta Ramasz, et al. Single-cell proteo-genomic reference maps of the hematopoietic system enable the purification and massive profiling of precisely defined cell states. Nature immunology, 22(12):1577–1589, 2021.

[14] Matthew T Witkowski, Igor Dolgalev, Nikki A Evensen, Chao Ma, Tiffany Chambers, Kathryn G Roberts, Sheetal Sreeram, Yuling Dai, Anastasia N Tikhonova, Audrey Lasry, et al. Extensive remodeling of the immune microenvironment in b cell acute lymphoblastic leukemia. Cancer cell, 37(6):867–882, 2020.

[15] Adam Gayoso, Zoë Steier, Romain Lopez, Jeffrey Regier, Kristopher L Nazor, Aaron Streets, and Nir Yosef. Joint probabilistic modeling of single-cell multi-omic data with totalvi. Nature methods, 18(3):272–282, 2021.

[16] Matthew P Mulè, Andrew J Martins, and John S Tsang. Normalizing and denoising protein expression data from droplet-based single cell profiling. Nature communications, 13(1):2099, 2022.

[17] Caibin Sheng, Rui Lopes, Gang Li, Sven Schuierer, Annick Waldt, Rachel Cuttat, Slavica Dimitrieva, Audrey Kauffmann, Eric Durand, Giorgio G Galli, et al. Probabilistic machine learning ensures accurate ambient denoising in droplet-based single-cell omics. bioRxiv, pages 2022–01, 2022.

[18] Yuan Yin, Masanao Yajima, and Joshua D Campbell. Characterization and decontamination of background noise in droplet-based single-cell protein expression data with DecontPro. Nucleic Acids Research, page gkad1032, 11 2023.

[19] Jeff A Bilmes et al. A gentle tutorial of the em algorithm and its application to parameter estimation for gaussian mixture and hidden markov models. International computer science institute, 4(510):126, 1998.

[20] Yuri Kotliarov, Rachel Sparks, Andrew J Martins, Matthew P Mulè, Yong Lu, Meghali Goswami, Lela Kardava, Romain Banchereau, Virginia Pascual, Angélique Biancotto, et al. Broad immune activation underlies shared set point signatures for vaccine responsiveness in healthy individuals and disease activity in patients with lupus. Nature Medicine, 26(4):618–629, 2020.

[21] Felix Sebastian Nettersheim, Sujit Silas Armstrong, Christopher Durant, Rafael Blanco-Dominguez, Payel Roy, Marco Orecchioni, Vasantika Suryawanshi, and Klaus Ley. Titration of 124 antibodies using cite-seq on human pbmcs. Scientific reports, 12(1):20817, 2022.

[22] Yuhan Hao, Stephanie Hao, Erica Andersen-Nissen, William M Mauck, Shiwei Zheng, Andrew Butler, Maddie J Lee, Aaron J Wilk, Charlotte Darby, Michael Zager, et al. Integrated analysis of multimodal single-cell data. Cell, 184(13):3573–3587, 2021.

[23] Diederik P Kingma and Max Welling. Auto-encoding variational bayes. arXiv preprint arXiv:1312.6114, 2013.

[24] Carl Doersch. Tutorial on variational autoencoders. arXiv preprint arXiv:1606.05908, 2016.

[25] David E Rumelhart, Geoffrey E Hinton, and Ronald J Williams. Learning representations by back-propagating errors. nature, 323(6088):533–536, 1986.

[26] Irina Higgins, Loic Matthey, Xavier Glorot, Arka Pal, Benigno Uria, Charles Blundell, Shakir Mohamed, and Alexander Lerchner. Early visual concept learning with unsupervised deep learning. arXiv preprint arXiv:1606.05579, 2016.

[27] Diederik P Kingma and Jimmy Ba. Adam: A method for stochastic optimization. arXiv preprint arXiv:1412.6980, 2014.

[28] Luca Scrucca, Michael Fop, T Brendan Murphy, and Adrian E Raftery. mclust 5: clustering, classification and density estimation using gaussian finite mixture models. The R journal, 8(1):289, 2016.

